# Linkage Disequilibrium and Population Structure in Wild and Cultivated Populations of Rubber Tree (*Hevea brasiliensis*)

**DOI:** 10.1101/360610

**Authors:** Livia Moura de Souza, Luciano H. B. dos Santos, João R. B. F. Rosa, Carla C. da Silva, Camila C. Mantello, André R. O. Conson, Erivaldo José Scaloppi, Josefino de Freitas Fialho, Mario Luiz Teixeira de Moraes, Paulo de S. Gonçalves, Gabriel R. A. Margarido, Antonio A. F. Garcia, Vincent Le Guen, Anete P. de Souza

**Author notes:** Both authors contributed equally to this work (Livia Moura de Souza, Luciano H. B. dos Santos). Corresponding Author: Anete Pereira de Souza.

## Abstract

Among rubber tree species, which belong to the *Hevea* genus of the Euphorbiaceae family, *Hevea brasiliensis* (Willd. ex Adr.de Juss.) Muell. Arg. is the main commercial source of natural rubber production worldwide. Knowledge of the population structure and linkage disequilibrium (LD) of this species is essential for the efficient organization and exploitation of genetic resources. Here, we obtained single-nucleotide polymorphisms (SNPs) using a genotyping-by-sequencing (GBS) approach and then employed the SNPs for the following objectives: (i) to identify the positions of SNPs on a genetic map of a segregating mapping population, (ii) to evaluate the population structure of a germplasm collection, and (iii) to detect patterns of LD decay among chromosomes for future genetic association studies in rubber tree. A total of 626 genotypes, including both germplasm accessions (368) and individuals from a genetic mapping population (254), were genotyped. A total of 77,660 and 21,283 SNPs were detected by GBS in the germplasm and mapping populations, respectively. The mapping population, which was previously mapped, was constructed with 1,062 markers, among which only 576 SNPs came from GBS, reducing the average interval between two adjacent markers to 4.4 cM. SNPs from GBS genotyping were used for the analysis of genetic structure and LD estimation in the germplasm accessions. Two groups, which largely corresponded to the cultivated and wild populations, were detected using STRUCTURE and via principal coordinate analysis (PCoA). LD analysis, also using the mapped SNPs, revealed that non-random associations varied along chromosomes, with regions of high LD interspersed with regions of low LD. Considering the length of the genetic map (4,693 cM) and the mean LD (0.49 for cultivated and 0.02 for wild populations), a large number of evenly spaced SNPs would be needed to perform genome-wide association studies in rubber tree, and the wilder the genotypes used, the more difficult the mapping saturation.

## 1 Introduction

*Hevea brasiliensis*, or the rubber tree, is an important crop species that produces a high-quality natural rubber in commercially viable quantity, accounting for more than 98% of the total natural rubber production worldwide (Priyadarshan and Goncalves, 2003). A native species of the Amazon rainforest, *H brasiliensis* is a diploid (2n=36, n=18), perennial, and cross-pollinated tree species with an estimated haploid genome size of 1.47 Gb (Tang *et al*., 2016). This species belongs to the *Euphorbiaceae* family, comprising 11 inter-crossable species, of which *H. brasiliensis* is the most economically important (Gonçalves et al. 1990). The rubber tree has a heterozygous nature, with a long growing cycle that includes 5 years before latex collection. Like most forest trees, the rubber tree has a long generation time, which explains the slow progress of breeding this species and elucidating the genetic architecture of its complex traits using traditional approaches. Genetic breeding programs are challenged by a low seed yield per pollination (an average of ten seeds obtained for 100 pollinated flowers) and inbreeding depression, making it difficult to develop the appropriate progeny for classical genetic studies (Lespinasse et al., 2000). Hence, relatively little is known about genome-wide models of recombination, allele frequency variation, and linkage disequilibrium (LD) in this important plant.

Over the last 15 years, many genetic maps of the rubber tree have been constructed. The first rubber tree marker-based genetic maps were built with restriction fragment length polymorphisms (RFLPs) and amplified fragment length polymorphisms (AFLPs) (Lespinasse *et al*., 2000), and dense genetic maps were subsequently constructed using simple sequence repeats (SSRs) (Le Guen *et al*., 2011; Triwitayakorn *et al*., 2011; Souza *et al*., 2013). Saturated genetic linkage maps are important for the identification of genomic regions containing major genes and quantitative trait loci (QTLs) controlling agronomic traits, and such maps are important for further breeding programs.

In recent years, advances in next-generation sequencing technology (NGS) have lowered the cost of DNA sequencing to the point that genotyping-by-sequencing (GBS) (Elshire *et al*., 2011) is now feasible for high-diversity, large-genome species, and a genetic map has been developed using the GBS approach (Pootakham *et al*., 2015). GBS utilizes restriction enzymes to capture a reduced representation of the target genome, and with DNA-barcoded adapters, it is possible to sequence multiple samples in parallel in a single run using an NGS platform. GBS has recently been applied to the large barley (*Hordeum vulgare* L.) and wheat (*Triticum aestivum*) genomes and has been shown to be an effective tool for developing molecular markers for these species (Poland *et al*., 2012).

Evaluation of the molecular diversity encompassed in rubber tree genetic resources is a prerequisite for their efficient exploitation in breeding and the development of conservation strategies of genetic diversity. de Souza *et al*. (2015) analyzed approximately one thousand cultivated rubber tree accessions originating from various geographic areas in a Brazilian germplasm collection. These accessions were genotyped with 13 SSR markers distributed across the chromosomes of the species, and a total of 408 alleles were identified, 319 of which were shared between groups, while 89 alleles were specific to different groups of accessions.

LD is the non-random association of alleles at distinct loci in the genome of a sampled population (Weir, 1979) and is the basis for association mapping approaches. LD can be used for many purposes in plant genomics research and has received considerable attention as a tool for the study of marker-trait associations due to physical linkage, followed by marker-assisted selection (MAS). Another important application of LD is the study of genetic diversity in natural populations and germplasm collections, where it can be employed for the evaluation of population genetics and in crop improvement programs, respectively (Gupta *et al*., 2005).

With the rise of sequence-based genotyping, precise and accurate estimates of population structure and the LD across the genome are now attainable for the rubber tree. Our goals in this study were to characterize the genome distribution of single-nucleotide polymorphisms (SNPs) in a rubber tree mapping population using GBS technology, to examine population structure, to investigate how LD breakdown relates to chromosomes, and to compare the LD between cultivated and wild populations. For this study, we used accessions selected from the germplasm collection previously analyzed by de Souza *et al*. (2015) and a mapping population (PR255 x PB217) previously saturated with SSR markers described by Souza *et al*. (2013) and Rosa *et al*. (submitted).

## 2 Material and methods

### 2.1 Plant materials and DNA extraction

Two sets of samples were selected for this study, and a total of 626 samples were sequenced. One set consisted of 368 *H. brasiliensis* accessions, composed of both wild germplasm and cultivated genotypes. Details of the plant materials can be found in de Souza *et al*. (2015) and Supplementary Table 1. The other set is an important mapping population of the rubber tree consisting of 252 F1 hybrids, derived from a cross between PR255 x PB217 and comprising three replicates of each parental genotype, which were mapped with 505 markers (SSRs, expressed sequence tag-SSRs, and SNPs) prior to this publication (Souza *et al*., 2013; Rosa *et al*., submitted). Genomic DNA was extracted from leaves using the DNeasy^®^ Plant Mini Kit (QIAGEN, Germany) according to the procedures described by the manufacturer. DNA quality parameters and concentrations were measured using a UV-Vis spectrophotometer (NanoDrop, Thermo Scientific, Wilmington, DE, USA) and agarose gels.

### 2.2 SNP discovery via GBS

GBS library preparation and sequencing were performed at the Institute of Genomic Diversity (Cornell University, Ithaca, NY, USA) as described by Elshire *et al*. (2011). Genome complexity was reduced by digesting individual genomic DNA samples with *EcoT22I*, a methylation-sensitive restriction enzyme. The resultant fragments from each sample were directly ligated to a pair of enzyme-specific adapters and combined into pools. PCR amplification was carried out to generate the GBS libraries, which were sequenced on the Illumina HiSeq 2500 platform (Illumina Inc., USA). The raw data were processed, and SNP calling was performed using TASSEL 5.0 (Glaubitz *et al*., 2014).

Initially, the FASTQ files were demultiplexed according to the assigned barcode. The reads from each sample were trimmed, and the tags were identified using the following parameters: Kmer length of 64 bp, minimum quality score within the barcode and read length of 20, minimum Kmer length of 20. All sequence tags from each sample were aligned to the reference rubber tree genome (Tang *et al*., 2016) with Bowtie 2 (Langmead and Salzberg, 2012) using the very-sensitive option.

To perform the analysis, the data were divided into the mapping population and germplasms. SNP calling was performed using the TASSEL 5 GBSv2 pipeline (Glaubitz *et al*., 2014) and filtered using VCFtools (Danecek *et al*., 2011) with the following criteria: (1) missing data of 20%, (2) minor allele frequency (MAF) greater than or equal to 5% (MAF 0.05), and (3) biallelic SNPs only.

### 2.3 Genetic linkage map

All linkage analyses were performed using OneMap software (Margarido *et al*., 2007), version 2.0-1, employing a previously constructed genetic map (Souza *et al*., 2013; Rosa *et al*., submitted) as a basis for the inclusion of GBS-based SNPs with a minimum logarithm of odds (LOD) score of 8.21 (according to the function in the R package Onemap “suggest_lod”) and a maximum recombination fraction of 0.35.

The map construction utilized only markers with 0.05 missing data and tested the pattern of allelic segregation for χ2 goodness of fit to the expected Mendelian segregation ratios, and markers with significant segregation distortion were excluded from further analysis. GBS-based SNPs were added to the previous genetic map using the ‘try.seq’ function in OneMap, which determines the best position for a given unpositioned GBS marker in a specific linkage group (LG). Finally, the fractions of recombination were converted to centimorgans (cM) using the Kosambi map function (Kosambi, 1943), and the map was drawn in Mapchart, version 2.3 (Voorrips, 2002).

### 2.4 Population structure and genetic diversity

The population structure was investigated in 368 genotypes from the germplasm collection with data from SNPs anchored in a certain LG using two different methods: STRUCTURE analysis and principal coordinate analysis (PCoA). Initially, the structure was analyzed with the software STRUCTURE 2.3.4 (Pritchard *et al*., 2000). Ten replications were run for each of the subpopulation numbers (K), ranging from 1 to 10. Each run included 500,000 Markov chain Monte Carlo (MCMC) iterations, among which the first 100,000 iterations (used to monitor whether a chain reached stationarity) were discarded as burn-in. The delta K method was used to identify the number of subgroups in the dataset (Evanno *et al*., 2005). Based on the posterior probability of membership (Q) of a given accession, the accession was classified as admixed in clusters with a membership of Q<0.70. Subsequently, genetic distances between pairs of accessions were calculated, and PCoA was performed for the SNPs using the GenAlEx program (version 6.5) (Peakall and Smouse, 2012).

### 2.5 Analysis of LD

LD was measured by calculating the squared correlation coefficient (*r*^2^) between each pair of SNPs with the R software and GGT 2.0 (van Berloo, 2008), using data from SNPs anchored in the genetic map. We selected markers that were positioned on the linkage map and were present in the germplasm to calculate the LD in each LG separately, considering the subgroup inferred with STRUCTURE. The decay of LD over genetic distance was investigated by plotting pair-wise *r^2^* values against the distance (cM) between markers on the same chromosome using the following model: *y* = *a* + *be ^−c/x^* (Ranc *et al*., 2012), where x and y represent the genetic distance in cM and the estimated *r*^2^, respectively. The critical *r*^2^ for LD decay was determined by values of 0.1, which is considered the minimum threshold for a significant association between pairs of loci and to describe the maximum genetic or physical distance at which LD is significant (Zhu *et al*., 2008).

## 3 Results

### 3.1 SNP discovery and evaluation

The analysis was performed separately for the mapping population and germplasm to produce a total of 1.785 million reads of sequence data, of which 89% (1,586 million reads) consisted of good barcoded reads. Of these reads, 69,408 (23.42%) were aligned to the mapping population exactly one time and 177,143 (59.78%) more than one time, corresponding to an 83.20% (246,551) overall alignment rate. The total rate of alignment to the germplasm was 89.97% (818,807), of which 22.59% of the tags (205,576) were aligned to the rubber tree reference genome exactly one time (Tang *et al*., 2016) and 67.38% (613,231) more than one time.

A total of 386,180 SNPs were identified in the germplasm, and 76,191 SNPs were detected in the mapping population, of which 350,965 SNPs (germplasm) and 66,453 SNPs (mapping population) were biallelic. After excluding markers showing (1) more than 20% missing data or (2) a MAF ≤ 0.05, the whole dataset was reduced to 77,660 and 21,283 SNPs in the germplasm and mapping populations, respectively. The SNP frequencies were one biallelic SNP every 20.7 kb for the mapping population and every 3.9 kb for the germplasm.

Sequence data are deposited under EMBL-EBI accession PRJEB26962.

### 3.2 Saturation of the linkage map with SNPs

From 21,283 SNPs, a total of 14,852 markers were selected after applying the chi-square test (*P* ≤ 0.05), which revealed segregation ratios of 1:2:1 and 1:1. Markers exhibiting statistically significant segregation distortion were excluded from further analysis to obtain accurate genetic linkage maps. The data analyses were performed using SNPs with a maximum of 5% missing data, resulting in a linkage map with 1,062 markers presenting 348 SSR markers (Souza *et al*., 2013 and Mantello, 2014), 576 SNPs from GBS (Supplementary Table 2) and 138 SNPs markers genotyped using the Sequenom MassARRAY^®^ platform (AgenaBio, San Diego, CA) and the Fluidigm^®^ platform (South San Francisco, CA), developed from *de novo* transcriptome assemblies (Mantello *et al*., 2014 and Salgado *et al*., 2014) and from EST full-length libraries (Silva *et al*., 2014).

The genetic map was organized according to the numbers obtained from the map previously developed by Rosa *et al*. (submitted). Eleven markers were removed after data diagnosis using heat map graphs, thereby permitting the visualization of the recombination fraction and LOD scores from markers, to group the SNPs in the LGs without changing the order of the base map. Only LG18 was further divided into subgroups “A” and “B” (Figure 1).

**Figure 1.**
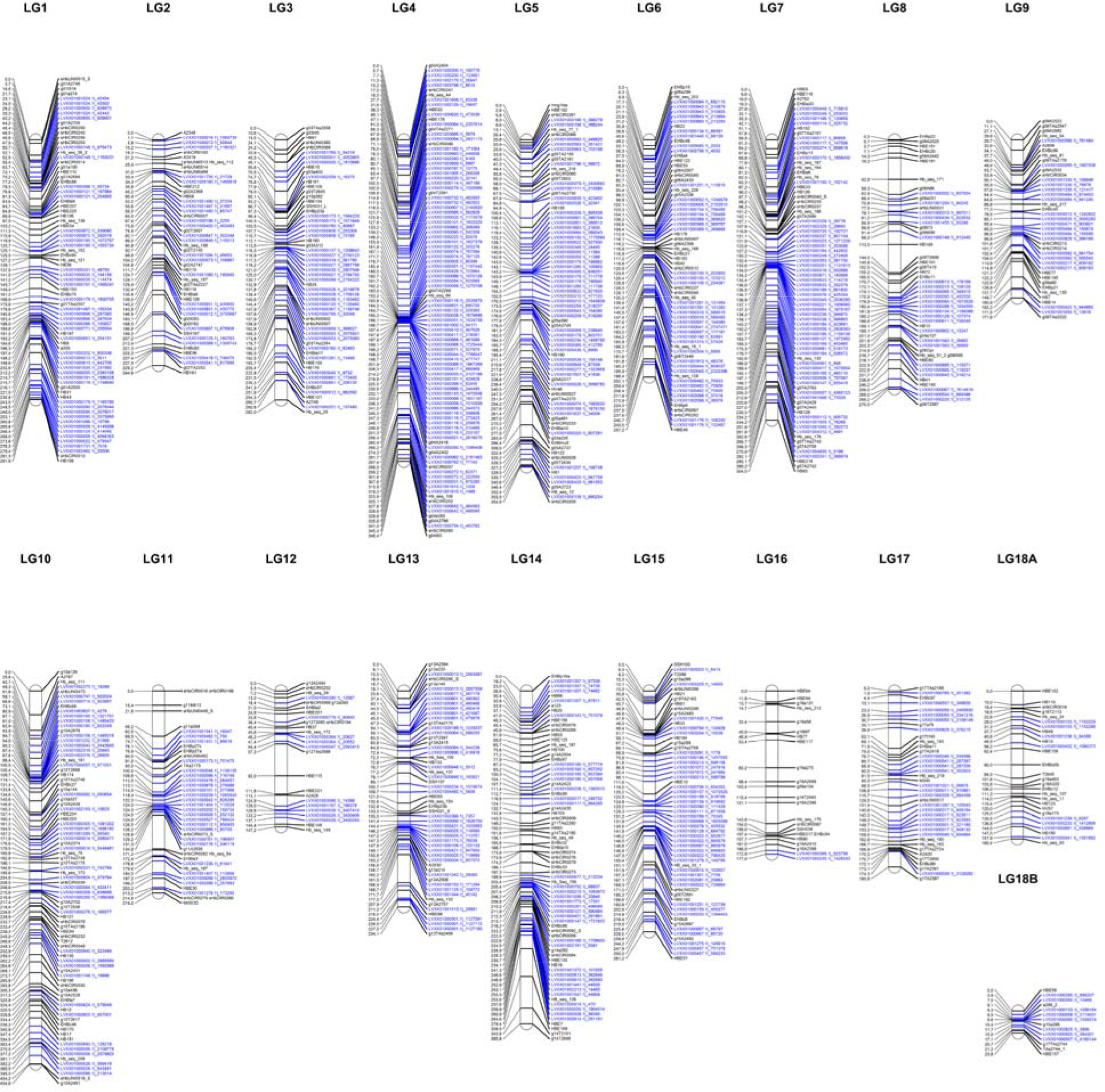
Linkage map of the rubber tree. Markers in blue represent SNPs obtained using the genotyping-by-sequencing technique, and markers in black were obtained from the previous map (Rosa *et al*., SUBMITTED).

Thus, a genetic map was generated spanning a cumulative length of 4,693 cM (Figure 1), distributed among all the chromosomes. The LGs ranged from 23.8 cM (LG18B) (Supplementary Table 3) with 14 markers to the largest group with 404.6 cM (LG10) with 85 markers (of which 78 were from GBS), and the average interval size between two adjacent markers was 4.4 cM.

The maximum gap size was 36.4 cM (LG11), which was maintained from the previous map. Furthermore, some regions could not be sampled using the selected restriction enzyme; in LG16, for example, only 2 GBS-based markers were added.

### 3.3 Genetic relationships among populations

The germplasm collection was selected in a prior work by Souza *et al*. (2015) examining the population structure with SSR markers. To confirm the population structure of the sample of selected individuals, we performed a new analysis as performed with the SNPs from GBS genotyping. Only mapped SNPs that were common in the germplasm collection were used. Clustering inference performed with *K* values ranging from 1 to 10 showed that the model likelihood increased steeply at *K*=2, followed by a drastic decline starting at K=3, suggesting that the optimal *K* value was 2. The assignment results for *K*=2 showed that some of the sampled individuals exhibited admixtures from two gene pools (Figure 2(A)): group 1 (red bars) mainly consisted of accessions originating from the Mato Grosso and cultivated genotypes, and group 2 (green bars) consisted entirely of wild accessions from Amazonas, Rondônia, Para, and Acre (identified as IS - Ilha Solteira). Twenty-one accessions showing a membership probability (Q value) below 0.70 were defined as admixed and were removed from subsequent analyses.

**Figure 2.**
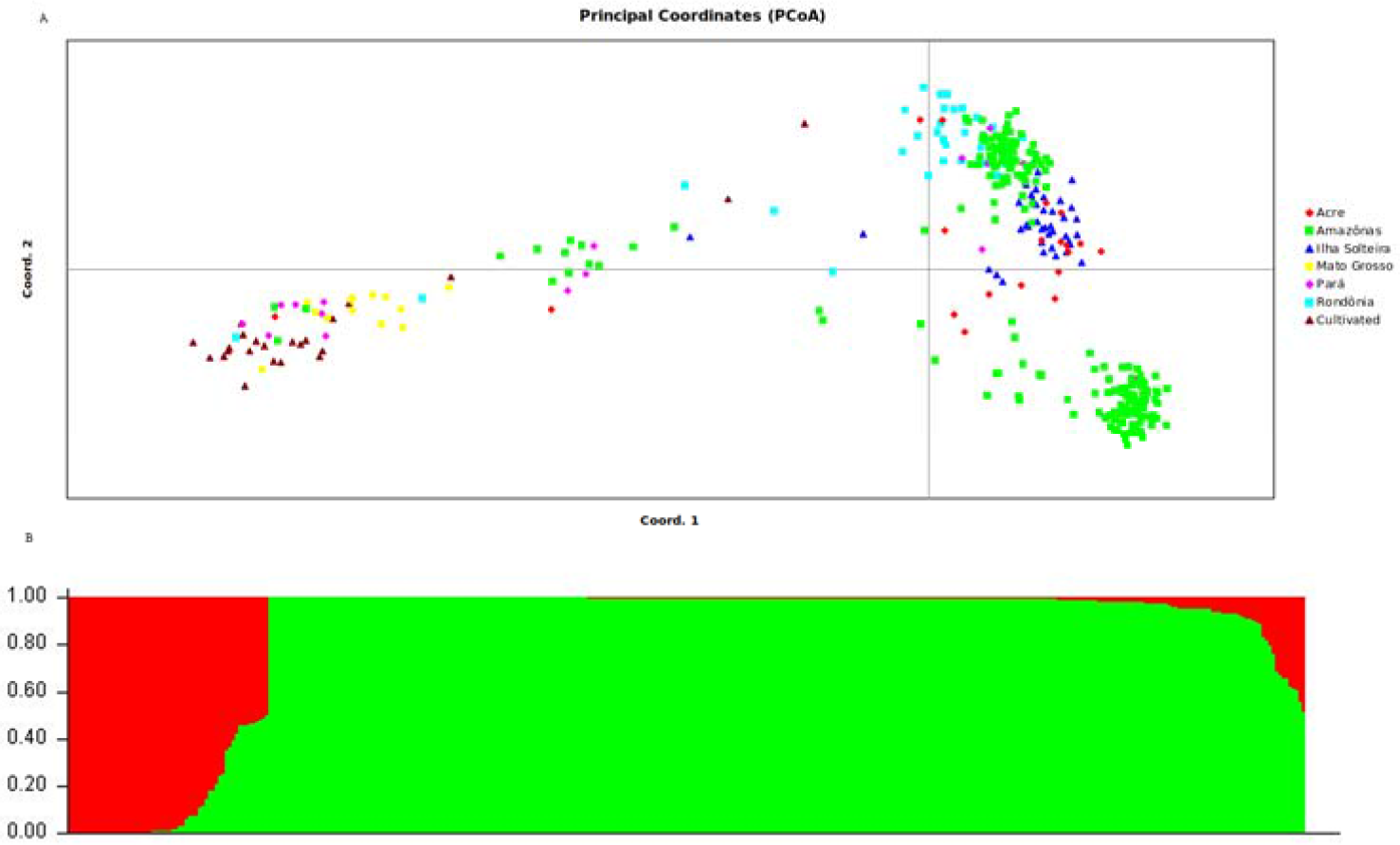
Estimated genetic structure of the wild population and breeding population of rubber tree based on PCoA (A) and STRUCTURE analysis (B). All analyses are based on the genetic variabilit of 438 SNP loci.

As a second analysis of differentiation, we employed PCoA based on a similarity matrix that explained 22.8% and 16.7% of the genetic variation with the first and second PCoA axes, respectively. Plotting the two first PCoA axes separated the germplasm into two clusters, though some overlap was present between the wild germplasms and cultivated genotypes (Figure 2). On the first axis, most of the breeding genotypes were separated from the other genotypes. On the second axis, the wild germplasm samples were clustered together, but some accessions from the Amazon were isolated in their own subdivision.

### 3.4 Evaluation of LD

Based on population genetic structure, accessions could be divided into two distinct groups (cultivated and wild group) (Figure 2), and pairwise LD estimates were performed within the gene pool of each of these groups. To visualize LD throughout the genome, heat maps were produced based on pairwise *r^2^* estimates, and corresponding *p-values* were calculated using permutations for all marker pairs (Supplementary Figure 1). These heat maps were employed to identify variations in LD between the cultivated and wild rubber tree germplasm groups.

In 16,025 pairwise combinations, we identified 186 (1.2%) and 592 (3.7%) statistically significant associations that were in LD (*P*<0.05) in the cultivated and wild germplasms, respectively. Of these significant associations, 78 and 64 were intrachromosomal in the cultivated and wild germplasms, respectively, accounting for 0.5% and 0.4% of the total possible intrachromosomal correlations. Among the unlinked loci, the proportions of LD were 0.7% and 3.3% for the cultivated and wild germplasms, respectively. In both clusters, an uneven distribution of LD among the 18 chromosomes was observed.

The strength of LD (*P*<0.05) was very different between two clusters, as reflected by the mean *r^2^* values of 0.49 and 0.02 obtained for the cultivated and wild germplasms, respectively. The 346 SNPs that were localized in the integrated map were used for the estimation of LD decay among the different LGs. LD decay was estimated across the germplasms and was found to be more pronounced in the wild germplasms, with the range being dependent on the chromosome group (Supplementary Figure 2). Using a fixed baseline *r^2^* value of 0.1 for the wild germplasm, LD decay ranged from 2.3 cM (LG9) to 11.4 cM (LG17). In contrast, in the cultivated accessions, this decay was slower, ranging from 4.3 cM (LG8) to 50 cM (LG17), with average values of 5.6 cM and 19.6 cM for the wild and cultivated germplasms, respectively (Figure 3). The patterns of LD can also be visualized across the genome from the diagonal of the heat maps (Supplementary Figure 1).

**Figure 3.**
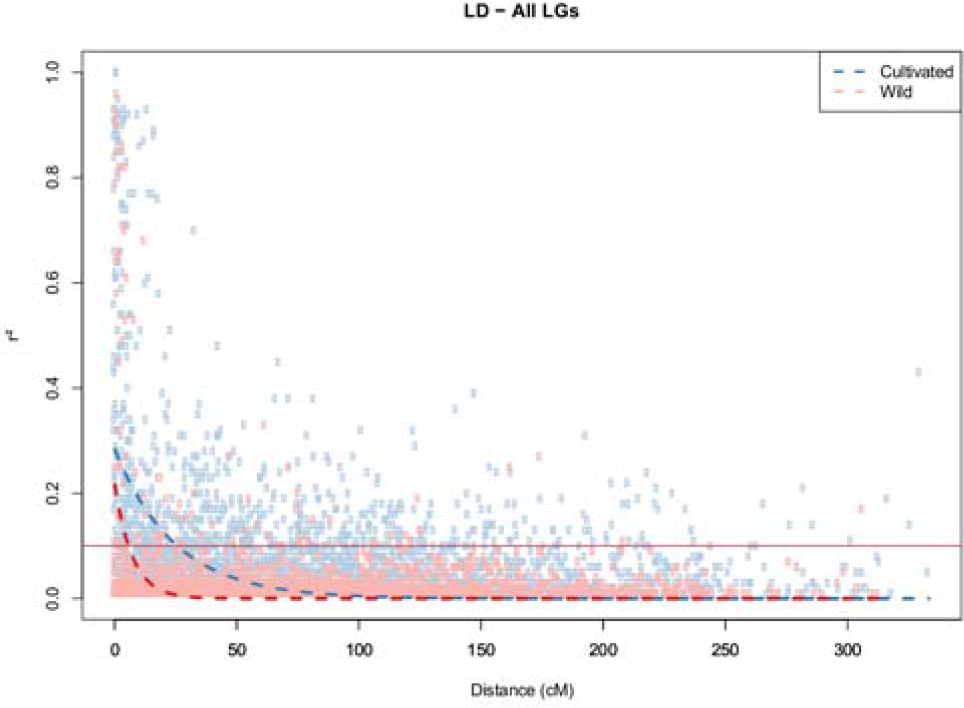
Decay of LD (r^2^) as a function of genetic distance (cM) between pairs of loci on all chromosomes. Only r^2^ values with *P*<0.05 are shown.

## 4 Discussion

We performed a genetic mapping study with a bi-parental population (252 segregating hybrids between PR255 x PB217) and analyzed the population structure and LD with two different populations of rubber tree, formed from 47 accessions from a breeding program and 300 accessions from a germplasm collection. For this study, we employed SNPs obtained via the GBS approach, which enabled the detection of polymorphisms distributed across the genome.

An important breakthrough of the GBS approach is that a reference genome is not necessary for SNP genotyping. However, the availability of a reference genome offers additional benefits, as it allows proper alignment and ordering of the sequenced tags (Poland *et al*., 2012). The first draft of the genome sequence (Rahman *et al*., 2013) provided a source of genomic information, after which three more genomes were published (Tang *et al*., 2016; Lau *et al*., 2016; Pootakham *et al*., 2017). The assembly is highly fragmented, containing more than one million contigs. The assembly of a complex genome is challenging, owing in part to the presence of highly repetitive DNA sequences, which introduce ambiguity during genome reconstruction. We used the genome published by Tang *et al*. (2016) to map the SNPs. The SNPs positioned on the map are named in reference to the location of alignment with the genome.

Repetitive regions account for 71% of the *Hevea* genome (Tang *et al*., 2016), posing a major challenge for the *de novo* assembly, particularly when exclusively short-read data are used. This phenomenon might explain the prevalence of tags that aligned to more than one unique region (59.78% to the mapping population and 67.38% to the germplasm) (Tang *et al*., 2016). Based on the properties of the reference genome (characterized by a great accumulation of repetitive sequences, primarily in heterochromatic regions) (Rahman *et al*., 2013), the restriction enzyme *EcoT22I* was selected because it is partially sensitive to methylation and rarely cuts retrotransposons.

The frequency of nucleotide substitutions was five times higher in the wild (3.9 kb) than in the cultivated (20.7 kb) germplasm sequences. One explanation for this finding is that selection in cultivated breeding programs acts to reduce diversity and alter allele frequencies in the DNA sequence. Depending on how LD is increased surrounding these loci; the effects of such a selection may not extend sufficiently far to affect the overall genome diversity. Rubber tree breeders have had to develop cultivars that are appropriate for the specific temperature and humidity conditions encountered in different cultivation areas, along with various biotic and abiotic stress resistance factors. High-density SNPs are also common in non-genetic regions where there is no selection pressure, and the abundance of these SNPs will be very useful for future assessments of breeding populations. The high diversity of SNPs is characteristic of outbreeding trees; in other studies examining rubber tree, Pootakham *et al*. (2011) previously identified a frequency of one SNP every 1.5 kb, while Pootakham *et al*. (2015) observed a frequency of one SNP every 308 nucleotides.

GBS does not require complete genome sequencing; only a targeted sequencing approach is necessary. Due to its high-throughput efficiency, GBS has been used for SNP identification and mapping in many plant species (Poland *et al*., 2012; Rabbi *et al*., 2014, Rimbert *et al*., 2018, Bekele *et al*. 2018). In rubber tree, most of the genetic linkage maps constructed to date have employed molecular markers such as RFLPs and AFLPs (Lespinasse *et al*., 2000) or microsatellites (SSRs) (Le Guen *et al*., 2011; Triwitayakorn *et al*., 2011; Souza *et al*., 2013). However, GBS was employed for linkage map construction in rubber tree very recently (Pootakham *et al*., 2015; Shearman *et al*., 2015). In the present study, we utilized the GBS platform to sequence a PR255 x PB217 mapping population that was previously saturated with SSR markers and SNPs obtained from other specific platforms (Souza *et al*., 2013; Rosa *et al*., submitted).

The linkage map constructed in the present study exhibits a regular marker distribution. However, the cumulative genetic map is significantly longer (4,693 cM) than would be expected for maps derived from other mapping populations that have presented a lower resolution; for example, a map of 2,052 cM (Pootakham *et al*., 2015) was obtained using only SNPs, and a map of 2,441 cM (Le Guen *et al*., 2011) was obtained using both SSRs and AFLPs. Such expansions of the rubber tree genetic linkage map have also reported previously (4,160 cM, Shearman *et al*., 2015). Several factors may be responsible for this phenomenon, including the numbers and types of mapped loci, the genetic constitution of different mapping populations and differences in mapping strategies, the mapping software and the ratio between the number of markers and the population size (Knox and Ellis, 2002).

Genetic mapping of the population (PR255 x PB217) was initiated in the first publication by Souza *et al*. (2013) using microsatellite markers. These first LGs were organized according to the numbers obtained from the map previously developed by Lespinasse *et al*. (2000), and information from other maps (unpublished) consisting of microsatellites in common was used to identify syntenic markers. Subsequently, the PR255 x PB217 mapping population was saturated with SSR markers and 243 SNPs obtained from platforms such as Sequenom MassARRAY iPLEX technology and KASP genotyping (Rosa *et al*., submitted). The SNPs from GBS used to construct the new genetic linkage map contributed to reducing the average interval between two adjacent markers (4.4 cM versus 7.4 cM). However, the marker density remained lower than that of the first rubber tree map (Lespinasse *et al*., 2000), which presented one marker every 3 cM or 0.89 cM according to Pootakham *et al*. (2015). LG11 displayed the maximum gap size (36.4 cM), which may have been caused by sections of the genome that were identical among the parental genotypes and thus an absence of polymorphisms, or by recombination hotspots. Large gaps exhibiting a low degree of polymorphisms have also been reported by Shearman *et al*. (2015).

Genetic maps are important tools not only for QTL mapping but also for anchoring genome assembly scaffolds into pseudo-chromosomes. The draft genome of *H. brasiliensis* has been reported to be highly heterozygous, with 71% of the genome length comprising repeats (Tang *et al*., 2016). A total of 21,283 biallelic SNPs were identified, potentially aligning with 1,926 scaffolds, corresponding to 26% of the total scaffolds of the genome used. However, only 3% of these scaffolds were anchored to LGs due to the relatively fragmented scaffold sequences and the usage of an insufficient genetic map to anchor the scaffolds. Thus, it is important to develop higher-quality genetic maps to help improve the scaffold anchoring ratio of future rubber tree genome assemblies.

Progress in the development of new molecular markers and genetic linkage maps is important for genetic improvement in rubber tree breeding programs. Despite the economic and ecological importance of this crop, restricted genomic resources are available for rubber tree. Sequencing of the studied family allowed the identification and genotyping of many markers in an efficient and cost-effective way. The linkage mapping analysis resulted in a number of LGs corresponding to the rubber tree haploid chromosome number (n=18) (Ong, 1975; Lespinasse *et al*., 2000), and the development of GBS methods and genetic maps represents an important advancement of the genomics tools available for these crops, which currently lack a good reference genome sequence.

Diversity analyses were performed in a previous work that included the entire collection of rubber germplasm, analyzing 1,117 genotypes with 13 microsatellite markers (Souza *et al*., 2015), and showed a mean observed heterozygosity (Ho) of 0.64 and higher genetic diversity (He) than Ho in all cases. Wright’s fixation index (F) values were positive, with a mean of 0.16 obtained for the accessions overall. Among a total of 408 observed alleles, 89 represented unique alleles to different groups of accessions, demonstrating the high level of heterogeneity in these genotypes. To confirm the population structure of the sample of selected individuals based on the work of de Souza *et al*. (2015), a new analysis of the population structure was conducted with SNPs obtained from GBS genotyping. The ΔK values obtained in this study indicated that the rubber tree germplasm could be divided into two groups and showed that some of the sampled individuals exhibited admixtures from two gene pools, which was also confirmed by plotting the two first PCoA axes.

Recent work using SSR markers has revealed similar results to those obtained using SNPs from GBS. For example, Le Guen *et al*. (2009) demonstrated a separation between the Acre and Rondônia groups and the Mato Grosso group. One possible reason for this separation is that the wild accessions are from geographically distant populations that are not connected by hydrographic networks (Le Guen *et al*., 2009; Chanroj *et al*., 2017), whereas most of the Wickham clones were collected from regions that are geographically closer to the Mato Grosso and the Tapajós river, enabling rubber seeds to flow from one region to another via the river. Souza *et al*. (2015) reported a difference in genotypes between the two groups, which are in different river basins, thus separating the genotypes in the cultivated and wild groups. Within the group of cultivated genotypes denoted as Mato Grosso and Wickhan, genotypes collected from Mato Grosso were genetically close to genotypes used to initiate the Asian breeding programs, denoted Wickhan genotypes in this article. These genotypes were collected by Henry Wickhan in 1976 in the same basin from which the genotypes from Mato Grosso were sampled (Gonçalves eta al., 1990). The results obtained by Souza *et al*. (2015) corroborate the seminal study by Le Guen *et al*. (2009).

Population genetic structure is a principal factor influencing the generation of false positives and the success of the association or LD mapping (Gupta *et al*., 2005). These 368 rubber tree accessions could be divided into two distinct groups based on population genetic structure: cultivated and wild germplasm. The investigation of LD decay *vs* genetic distance based on markers is not possible without prior information regarding the positions of markers in the genome. Since the information on the rubber tree genome is still inaccurate regarding these positions, we employed the existing genetic map of the PR255 x PB217 population to identify the positions of some SNPs and, thus, enable a more accurate study of LD decay.

LD analysis with the mapped SNPs revealed that LD varied along the chromosomes, with regions of high LD being interspersed with regions of low LD (Supplementary Figure 1). LD estimation is possible without the positions of molecular markers along the genome and was indeed performed in this study. However, these positions are crucial if one is interested in determining the decay or extension of LD in relation to the genetic distance, regardless of by chromosome.

The average *r^2^* values were found to be very different when the two detected groups were compared. This measurement was higher in the breeding germplasm (0.49) than in the wild germplasm (0.02), corroborating the results of Chanroj *et al*. (2017), who suggested that these results were caused by high gene flow in the wild Amazonian population. The breeding accessions exhibited a notably higher level of LD, suggesting less genetic diversity in this subdivision, perhaps because of the constraint of genetic variability employed in breeding programs due to the recurrent process of selection. The low LD detected in the wild germplasm in this study was expected because perennial outcrossing tree species display a high effective recombination rate, which leads to the rapid decay of LD (Krutovsky and Neale, 2005).

The higher LD level detected in the cultivated group may have been influenced by the partly identical-by-descent of these genotypes from a limited number of founders of the breeding programs, with only a few generations between them. Thus, many longer pieces of chromosomes have not had time to undergo disruptions. Domesticated crop cultivars necessarily represent a subset of the genetic variation found in their wild ancestors, and the process of crop domestication is responsible for genetic bottlenecks (McCouch, 2004). Although rubber tree breeding programs are very recent, the differences in LD patterns detected between cultivated and wild germplasms suggest that plant breeders may have selected for separate combinations of genes during the breeding process. Moreover, selection for high latex yields and the extensive use of particular clones as parents in rubber breeding programs have further reduced the genetic diversity of commercial rubber germplasm (Priyadarshan, 2016), which may also affect LD.

LD decay was estimated at 25.7 cM within cultivated and 5.7 cM within wild germplasm, and significant inter-chromosomal LD was identified within cultivated in contrast to wild germplasm. The distances of LD decay between the two groups were most different in LGs 3, 5, 7, 10 and 11, suggesting that these chromosomes may carry more genes related to agronomic traits that have been selected via organized breeding of this crop. In previous studies involving Amazonian accessions of rubber tree, Chanroj *et al*. (2017) revealed an LD decay of more than 0.5–6 cM and suggested that LD estimates were significantly influenced by physical distance, with LD decay greater than 2 kbp being observed in the widespread Amazonian population. These authors showed that LD decay over genetic distance was different for the 18 different chromosomes, possibly because of the different recombination rates of the 18 chromosomes. Rapid LD decay has been reported for many other outcrossing tree species, such as *Populus nigra*, in which a decay of *r^2^* with distance in the *CAD4* gene was observed at approximately 16 bp (Marroni *et al*., 2011). In *Eucalyptus globulus*, candidate genes for wood quality were analyzed using SNPs, and LD was estimated to decay rapidly (Thavamanikumar *et al*., 2011). Most LD estimation studies conducted in tree species are based on candidate genes (Krutovsky and Neale, 2005).

LD mapping relies on germplasm samples and, as such, does not require the development of experimental crosses with specific genetic backgrounds, and thus ease of use is an obvious benefit in studies of perennial species with long life cycles. LD is a key factor in determining the number of markers needed for genome-wide association studies (GWAS) and genomic selection (GS). Genomes with low LD require a high marker density for GWAS or GS; therefore, our SNPs may be valuable for GWAS in rubber tree breeding. Considering the length of the genetic map (4,693 cM) and the mean LD observed (0.49 in breeding and 0.02 in wild populations), many evenly spaced SNPs would be necessary to perform GWAS in the rubber tree, and the wilder the genotypes that are used, the more difficult is the saturation of the mapping. However, to obtain a sufficient SNP density throughout the genome and to account for variation in LD along the chromosomes more markers must be genotyped. Our study results provide a valuable resource for further genetic studies involving linkage or association mapping, marker-assisted breeding and *Hevea* sequence assembly and comparative mapping.

Furthermore, GWAS of wild germplasm accessions in the future will provide a substantial contribution to dealing with new challenging situations that will arise as a consequence of global climatic changes. Useful QTLs and genes to face these new situations are currently unknown, but the best way to identify them would be through analyses relying on GWAS of large panels of wild genotypes. A precise assessment of the LD pattern across the genome of *Hevea* is necessary for such endeavors, and the present study supplies a major contribution to this goal.

## Author contributions

LMS, LHS, VG and AS designed the study and performed the experiments; LMS, LHS, AC, CS, CM, EJ, JF, and MM performed the experiments; LMS, LHS, JR, GM, VG, MM, PG and AG analyzed the data; and LMS, LHS and AS wrote the manuscript.

---

**Supplementary Figure 1.**
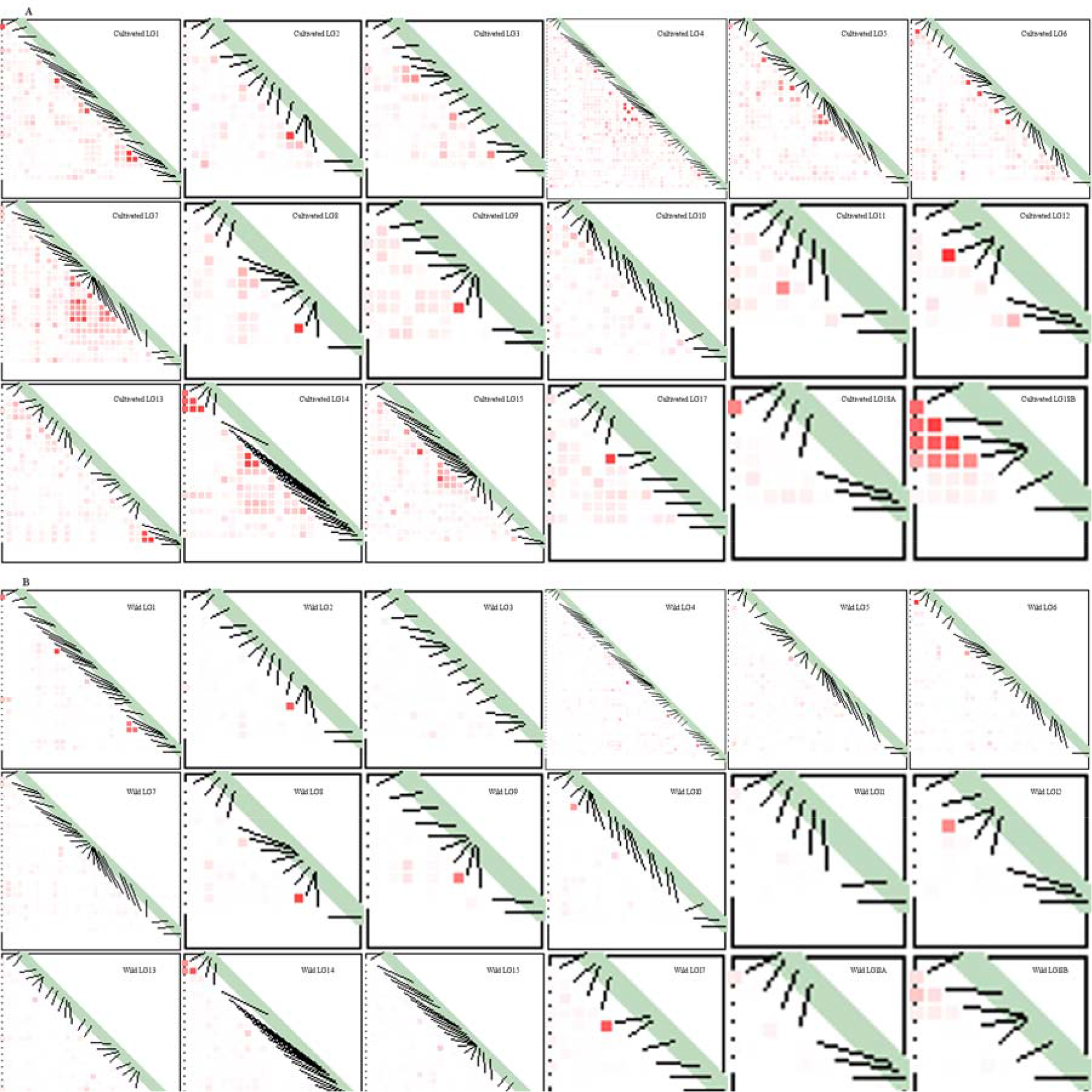
Plots of LD heat maps for (A) with 47 breeding accessions and (B) with 300 accessions from wild germplasms. The rubber tree LGs are represented by a diagonal bar. Markers were ordered on the x- and y-axes based on genomic location; therefore, each cell of the heat map represents a single marker pair. The r^2^ values for each marker pair are presented in the bottom half of the heat map and are represented by shades of red increasing in intensity in equal increments of 0.1 from 0.0 (white) to 1.0 (red).

**Supplementary Figure 2.**
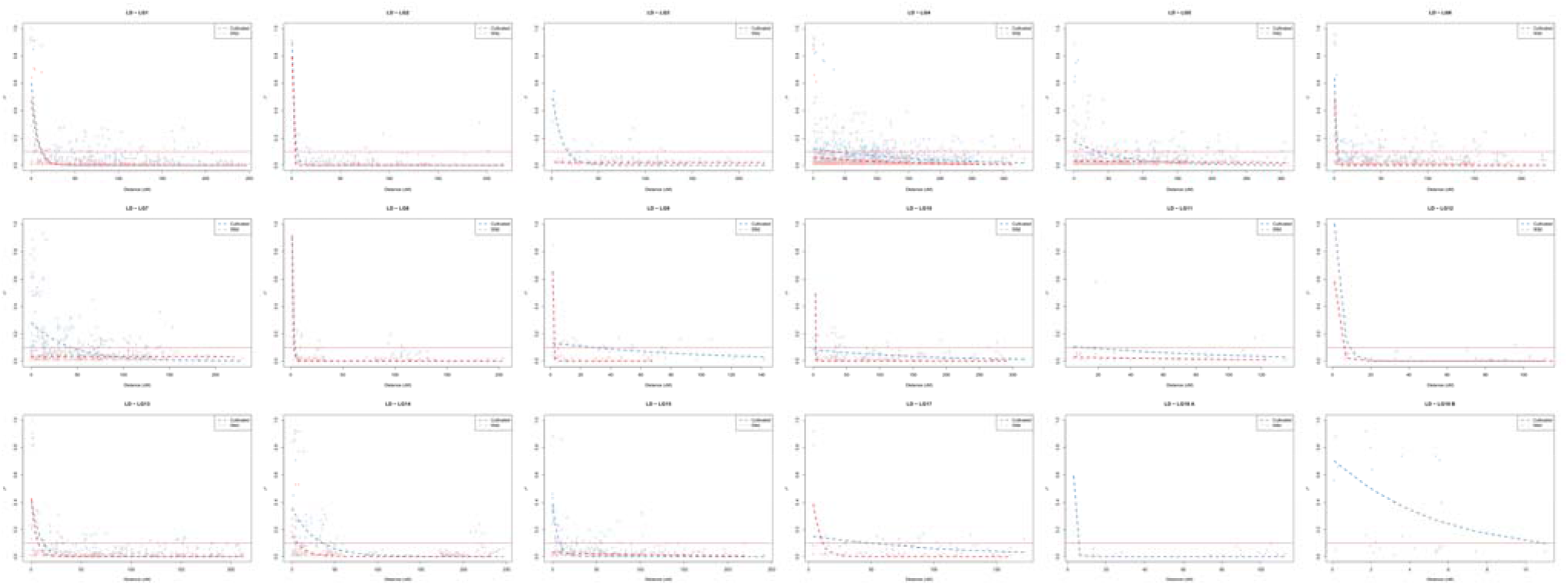
Decay of LD (r^2^) as a function of genetic distance (cM) between pairs of loci in individual LGs. Only r^2^ values with *P*<0.05 are shown.

**Supplementary Table 1.** Origin of germplasm genotypes and population structure results.

**Supplementary Table 2.** SNPs from GBS and their genome information. The name of markers are in agreement with the sequences of the reference genome (Tang *et al*., 2016).

**Supplementary Table 3.** Marker information for the genetic map.

